# Large-scale intrinsic connectivity is consistent across varying task demands

**DOI:** 10.1101/407205

**Authors:** Paulina Kieliba, Sasidhar Madugula, Nicola Filippini, Eugene P. Duff, Tamar R. Makin

**Affiliations:** Institute of Cognitive Neuroscience, University College London, 17 Queen Square, London WC1N 3AZ, UK; FMRIB Centre, University of Oxford, Oxford OX3 9DU, United Kingdom; Oxford Centre for Human Brain Activity, Wellcome Centre for Integrative Neuroimaging, Department of Psychiatry, University of Oxford, UK

**Keywords:** functional connectivity, resting state, steady-states, fMRI, ICA

## Abstract

Measuring whole-brain functional connectivity patterns based on task-free (‘restingstate’) spontaneous fluctuations in the functional MRI (fMRI) signal is a standard approach to probing habitual brain states, independent of task-specific context. This view is supported by spatial correspondence between task- and rest-derived connectivity networks. Yet, it remains unclear whether intrinsic connectivity observed in a resting-state acquisitions is persistent during task. Here, we sought to determine how changes in ongoing brain activation, elicited by task performance, impact the integrity of whole-brain functional connectivity patterns. We employed a ‘steady-states’ paradigm, in which participants continuously executed a specific task (without baseline periods). Participants underwent separate task-based (visual, motor and visuomotor) or task-free (resting) steady-state scans, each performed over a 5-minute period. This unique design allowed us to apply a set of traditional resting-state analyses to various task-states. In addition, a classical fMRI block-design was employed to identify individualized brain activation patterns for each task, allowing to characterize how differing activation patterns across the steady-states impact whole-brain intrinsic connectivity patterns. By examining correlations across segregated brain regions (nodes) and the whole brain (using independent component analysis), we show that the whole-brain network architecture characteristic of the resting-state is robustly preserved across different steady-task states, despite striking inter-task changes in brain activation (signal amplitude). Subtler changes in functional connectivity were detected locally, within the active networks. Together, we show that intrinsic connectivity underlying the canonical resting-state networks is relatively stable even when participants are engaged in different tasks and is not limited to the resting-state.

**New and Noteworthy**

Does intrinsic functional connectivity (FC) reflect the canonical or transient state of the brain? We tested the consistency of the intrinsic connectivity networks across different task-conditions. We show that despite local changes in connectivity, at the whole-brain level there is little modulation in FC patterns, despite profound and large-scale activation changes. We therefore conclude that intrinsic FC largely reflects the a priori habitual state of the brain, independent of the specific cognitive context.

## Introduction

Functional connectivity (FC) is a powerful and widely used tool for probing brain network organization and function in healthy (Biswal et al. 2010; Fair et al. 2007; Fox and Raichle 2007; Greicius and Menon 2004; Van Dijk et al. 2010) and clinical populations (Eippert et al. 2017; Filippini et al. 2009; Fox and Greicius 2010; Gilaie-Dotan et al. 2013; Hahamy et al. 2015a; Hahamy et al. 2015b). Many studies focus on FC measured during rest, which can be accurately described by a relatively small number of spatiotemporal patterns that remain consistent across different participants and datasets (Damoiseaux et al. 2006). The spatial composition of these resting-state patterns, often referred to as intrinsic connectivity networks (ICNs), have been shown to mirror the respective brain states during task execution (Gilaie-Dotan et al. 2013; Hahamy et al. 2017; Smith et al. 2009; Tavor et al. 2016; Wilf et al. 2017).

The high correspondence between rest- and task-based FC patterns (Cole et al. 2014; Fox et al. 2006; Greicius and Menon 2004; Moeller et al. 2009; Smith et al. 2009), as well as the changes in FC patterns between neurotypical and abnormal individuals, have led researchers to suggest that resting-state FC reflects the underlying synaptic efficacies in cortical networks (Guerra-Carrillo et al. 2014; Harmelech and Malach 2013; Kelly and Castellanos 2014; Sadaghiani and Kleinschmidt 2013). That is, intrinsic FC is suggested to reflect the habitual state of the brain, independent of the specific context. However, many new studies, such as those employing psychophysiological interactions (PPI; O’Reilly et al. 2012), emphasize the differences in FC patterns resulting from dynamic changes in task demands (Buckner et al. 2013; Hermundstad et al. 2013; Mennes et al. 2013; Shirer et al. 2012; Spadone et al. 2015). These divergent observations raise the question of whether FC, as measured using fMRI, is sensitive to changes in brain activation. Does intrinsic FC reflect the canonical (default, activation-independent), or current (transient, activation-dependent) state of the brain?

In this study, we sought to shed some light on this matter by testing the consistency of the ICN patterns across various well-defined steady-state tasks. We hypothesized that if intrinsic FC represents the canonical state of functional brain organization (i.e. synaptic efficacy; Harmelech and Malach 2013), it should remain relatively stable across changing tasks. Alternatively, if fMRI FC represented the transient task-dependent organization of the brain (Buckner et al. 2013; Hermundstad et al. 2013), intrinsic connectivity would be expected to change, depending on activation changes within the networks. We analyzed four steady-state conditions, collected either during rest or during three continuous tasks (without rest periods), allowing us to make inferences about resting and task-derived FC patterns based on the entire scan, rather than on brief rest and task periods, used in traditional block designs. This unique design also allowed us to employ multiple fMRI analyses developed for studying resting-state FC (otherwise not suitable for the more standard block-design), to study task-state FC and minimized the influence of rest on the task-based ICNs. Note that steady-state scans have been previously shown to be less susceptible to confounding factors than block designs (Hampson et al. 2006) and to produce more consistent FC results (Fair et al. 2007).

Previous studies have highlighted high degree of spatial overlap between rest and task-derived FC networks (Smith et al. 2009). Here we took a step further and interrogated connectivity strength between distinct brain regions to determine whether intrinsic connectivity remains stable across both task and rest. The different active steady-states used in our study were chosen based on a factorial design (motor/visual on/off) and were designed to target well-characterized and robust activation profiles in distinct sets of brain areas, with high consistency within and across participants. The natural vision condition was designed to activate the entirety of the Occipital- and Lateral Visual canonical ICNs. The motor task was performed with the right hand and was thus designed to only activate the left sensorimotor cortex, which comprised a portion of the bilateral sensorimotor ICN. The visuomotor condition was designed to simultaneously activate both visual and motor nodes. In addition to the 5-minute steady-state scans, we also used a traditional task-activation localizer (30 second blocks interleaved with baseline periods, see further details in the Methods) to measure changes in mean brain activation (BOLD signal level) induced by each of the tasks employed in the steady-state scans, and in each participant, allowing for participant-specific customized analysis.

We first used these data to determine the relationship between steady-state task-induced activation (based on the localizer task) and the FC profile (based on the various steady state scans) across brain nodes. We then employed a data driven approach based on independent components analysis (ICA) to investigate the stability of the ICNs, given the changed input induced by each steady-state task. For this purpose, we utilized the resting-state dataset collected by the Human Connectome Project as a model of the resting state ICNs (Smith et al. 2013). Both of these analyses showed little modulation in whole-brain FC patterns based on changed task activation. Finally, we used dual-regression analysis (Filippini et al. 2009) to isolate network-specific local changes in connectivity. By demonstrating that the overall architecture of the ICNs is highly robust despite changing task demands and specific, localized changes in FC, we conclude that intrinsic FC largely reflects the a priori habitual state of the brain, independent of the specific context.

## Methods

### Participants and Experimental Design

15 healthy volunteers (7 females, 8 males, age=27.25±4.4yr, all right handed) without any previous neurological disorders participated in the study after providing written informed consent. Participants were recruited in accordance with NHS national research ethics service approval (10/H0707/29). All participants underwent steady-state fMRI along with task localizer fMRI, with the order of the scans randomly determined. One participant was discarded from final analysis, due to an error in the block design acquisition. We note that the group ICNs in our dataset were comparable to those obtain using large datasets (HCP, see below), indicating adequate statistical power for the FC analysis. This dataset has been previously used to investigate related research questions (Costa et al. 2015; Duff et al. 2013; Duff et al. 2017).

Participants were scanned under four separate, five-minute continuous steady-state conditions (with no baseline epochs): rest, motor only, visual only, and simultaneous (but independent) visual and motor tasks (Figure 1A). The motor condition involved continuous sequential finger tapping against the thumb, using the right hand. Participants were asked to maintain a tapping frequency of 1Hz, and tapping pace was practiced prior to the scan. The natural vision condition consisted of videos of colorful abstract shapes in motion (Supplementary Video 1), modified from the work of the artist Len Lyn (circa 1930’s). During the combined visuomotor condition participants viewed the aforementioned videos while simultaneously performing the self-paced motor tapping task. The usage of ecologically-valid “low-level” tasks allowed us to construct a factorial design for activation profiles (i.e. orthogonal/additive activation in visual and motor areas across steady-states). In an additional fifth scan, described in the Supporting Information section, participants repeated the combined visuomotor condition, but were asked to change the finger-tapping direction (index-to-pinkie and reverse) whenever they noticed monochrome frames inserted in the video. This task was designed to explore the role of attentional load on the ICNs’ integrity. The order of scans was counterbalanced across participants, such that different participants were presented with the various scans using different, but complementary, order. A fixation cross was presented in all conditions and participants were asked to keep their eyes on the cross throughout the study.

**Fig. 1.**
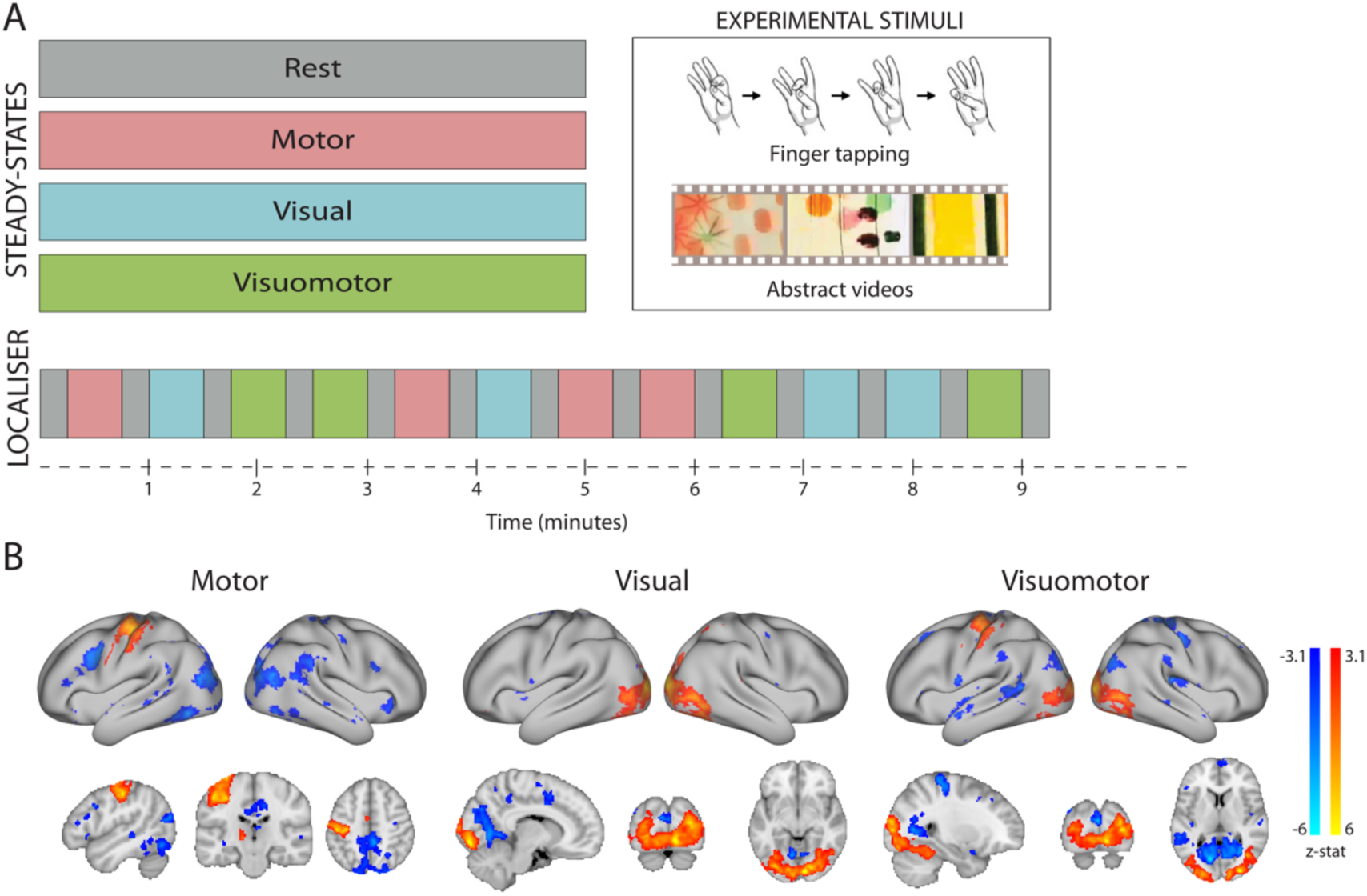
Steady-states design and localizer of activated brain areas. (A) Participants underwent a set of separate task-based (visual, motor and visuomotor) or task-free (resting) steady-state scans, each acquisition lasting five minutes. In addition, a task-localizer scan was employed to identify brain activation induced by each of the steady-state tasks. (B) Group activation maps during each of the steady-states employed in the localizer scan. All activation foci are projected onto the inflated surface of a template brain as well as on three anatomical planes.

An additional task-activation localizer scan was performed under the same conditions to identify participant-specific changes in average activation levels during each of the tasks. This scan used pseudo-randomized block design consisting of 30 second task intervals separated by 15 second baseline periods. During the localizer scan, each of the three main study tasks (visual only, motor only, visuomotor) was repeated four times for a total scan time of 9 minutes and 15 seconds (Fig. 1A).

### Data Acquisition

Functional data were acquired in a Siemens Vario 3T scanner, using a 32-channel head coil and a high-resolution multiband (factor 6) sequence with the following parameters: voxel size=2mm isotropic, TR=1300ms, TE=40ms, flip angle=66° (Feinberg et al. 2010; Moeller et al. 2010). 72 slices with 2mm thickness and no slice gap were acquired in the oblique axial plane, covering the whole cortex and cerebellum. Total number of volumes acquired: 230. For the task-localizer, the blood-oxygenation level dependent (BOLD) fMRI signal was acquired using a multiple gradient echo-planar T2*-weighted pulse sequence, with the parameters: voxel size= 3mm isotropic, TR=3000ms; TE=30ms; flip angle=90°; imaging matrix=64×64; FOV=192mm axial slices. Forty-six slices with slice thickness of 3mm and no gap were acquired in the oblique axial plane, covering the whole cortex, with partial coverage of the cerebellum. Total number of volumes acquired: 185. Anatomical data were acquired using a T1-weighted magnetization prepared rapid acquisition gradient echo sequence (MPRAGE) with parameters: TR: 2040ms; TE: 4.7ms; flip angle 8°; 1mm isotropic resolution. Field maps were obtained in order to reduce spatial distortion of the EPI images.

### Data pre-processing

All imaging data were processed using FSL-FEAT (version 6.00; Smith, 2004). Preprocessing included motion correction, field-map correction (Jenkinson et al. 2002) and brain extraction (Smith, 2002). Localizer scans only were subjected to spatial smoothing using a Gaussian kernel of FWHM of 5mm. Following the Human Connectome Project’s (HCP, http://www.humanconnectomeproject.org) minimal processing protocol, no spatial smoothing was applied to the steady-states data (Glasser et al. 2013). To account for the influence of any non-neuronal contribution to the BOLD signal, steady-state data were additionally cleaned using FIX (FMRIB’s ICA-based Xnoiseifier) (Griffanti et al. 2014) automated denoising. EPI volumes were spatially realigned to the mean image and co-registered with the structural T1weighted image using Boundary-Based Registration (Greve and Fischl, 2009). Time-course pre-whitening was carried out using FILM (FMRIB’s Improved Linear Model) with local autocorrelation correction (Smith et al. 2004). All structural and functional images were registered to standard MNI using both linear (FLIRT) and non-linear (FNIRT) registration. Images underwent mean-based intensity normalization and high-pass temporal filtering (0.01Hz for steady-state scans; 0.005Hz for localizer scans).

### Task localizer

A multi-level general linear model (GLM) analysis of the pre-processed localizer scans was used to identify regions that activated during one or more of the conditions, relative to rest (Jenkinson et al. 2012). The block design time-course of each of the four steady-state conditions was convolved with the gamma function, and together with its temporal derivative used to model the activation time-course in individual participants. Two participant-level contrasts were defined for each of the task conditions (motor, visual, visuomotor) vs rest (task>rest and rest>task), resulting in a total of 6 contrasts (2 per each task condition). A high-level group analysis was performed using a mixed effects model (Figure 1B). Z (Gaussianised T/F) statistic images were thresholded using clusters determined by Z>3.1 and a family-wise-error corrected cluster significance threshold of p<0.05 was applied to the suprathreshold clusters.

To verify that the regions activated during the task localizer were also activated during the steady-state scans we performed a region of interest (ROI) analysis, driven by the task-localizer results. The group task-localizer maps from motor, visual and visuomotor (task>rest) task conditions were thresholded (Z>3.1), binarized and used as the ROIs. The steady-states time-courses of the rest, motor, visual and visuomotor conditions were extracted from under the corresponding task-based ROI separately for each participant. The average amplitude of the time-course was compared between the rest and task-steady states using paired t-tests. Note that here a different pre-processing procedure was applied, including head-motion correction, field map correction, brain extraction, high-pass filtering, and spatial smoothing with 5 FWHM kernel.

### Node parcellation and network matrix generation

To measure inter-regional FC changes across the entire brain in each steady-state condition, an automatic brain parcellation was used. We used a multi-modal surface-based parcellation provided within the HCP, including 180 cortical regions in each hemisphere (Glasser et al. 2016). To generate participant- and condition-specific parcellations, the HCP parcellations were further dissected based on individual participants’ thresholded (Z>2.3) and binarized localizer scans (task>baseline). In other words, if the HCP ROI was found to be partially overlapping with a task activation cluster, that ROI was split into two separate nodes, each containing only activated/not activated voxels. This allowed us to generate nodes that are either task-relevant or task-irrelevant for each participant and condition (motor, visual, visuomotor). Functional connectivity matrices were then created by correlating the average time-course of each ROI with the average time-course of each of the other ROIs, separately for each individual participant and for each condition. Defining the nodes individually for each participant (based on their activation maps in conjugation with HCP parcellation) allowed us to take into account the inter-participant functional variability. Note that the activation maps used for node definition (task-localizer) were not used in the node analysis, therefore avoiding circularity.

Next, we investigated whether changes in correlations between nodes during rest and each of the steady-state tasks were associated with local changes in task-related activation. Specifically, we tested whether the changes in FC between a pair of nodes were associated with one or both of those nodes being activated (Zuo et al. 2018). First, to classify nodes based on their level of task-activation the following criteria ware applied: A single node was classified as task-relevant (activated) if, based on a given individual participant’s task-localizer (task>baseline), the average z-statistic value (across all voxels within that node) was above 2.3. A node was labelled as task irrelevant (not activated) if, based on the same criterion, the mean z-statistic value was higher than −1 and lower than 1. Nodes producing values outside these criteria (including the deactivated nodes) were discarded from this analysis (Zuo et al. 2018). Note that the deactivated nodes were not analyzed as, to date, the physiological interpretations of negative BOLD modulations remain controversial (Shih et al. 2009; Bianciardi et al. 2011; Hu & Huang 2015). Below we report results involving all of the study participants. Note, however, that the same pattern of results was observed when excluding participants with low number of activated nodes in a given condition (if their number of activated nodes was lower than 1.5 interquartile range (IQR) below the lower quartile (Q1) of number of activated nodes across all participants).

Next, the correlation coefficients across each of the steady-state time-courses were calculated across all pair-combinations of the nodes. Each pair of nodes was then sorted based on whether both nodes, one node, or neither of the nodes were activated during the visual, motor or visuomotor conditions. The correlation values across those node-pairs were Fisher’s z-transformed and displayed in a histogram, separately for each participant, task (visual, motor and visuomotor) and number of activated nodes (no nodes activated/one node activated/two nodes activated, see Fig. 2A and Supp. Fig. 1-8). Within each pair-category (none/one/two nodes activated) the average correlation coefficient (Fisher’s z-transformed) was further calculated for each participant and condition, and the mean values were analyzed using repeated-measures ANOVA and Bayesian repeated-measures ANOVA with number of task-related nodes in the pairwise correlation (none/one/two) and steady-state task (task/rest) as within-subject factors. This analysis was carried out in JASP (The JASP Team 2017), separately for motor, visual and visuomotor conditions. We note that averaging the FC across nodes may potentially mask subtler connectivity changes within each node category, which was the focus of an additional specialized analysis (dual regression approach, see below).

**Fig. 2.**
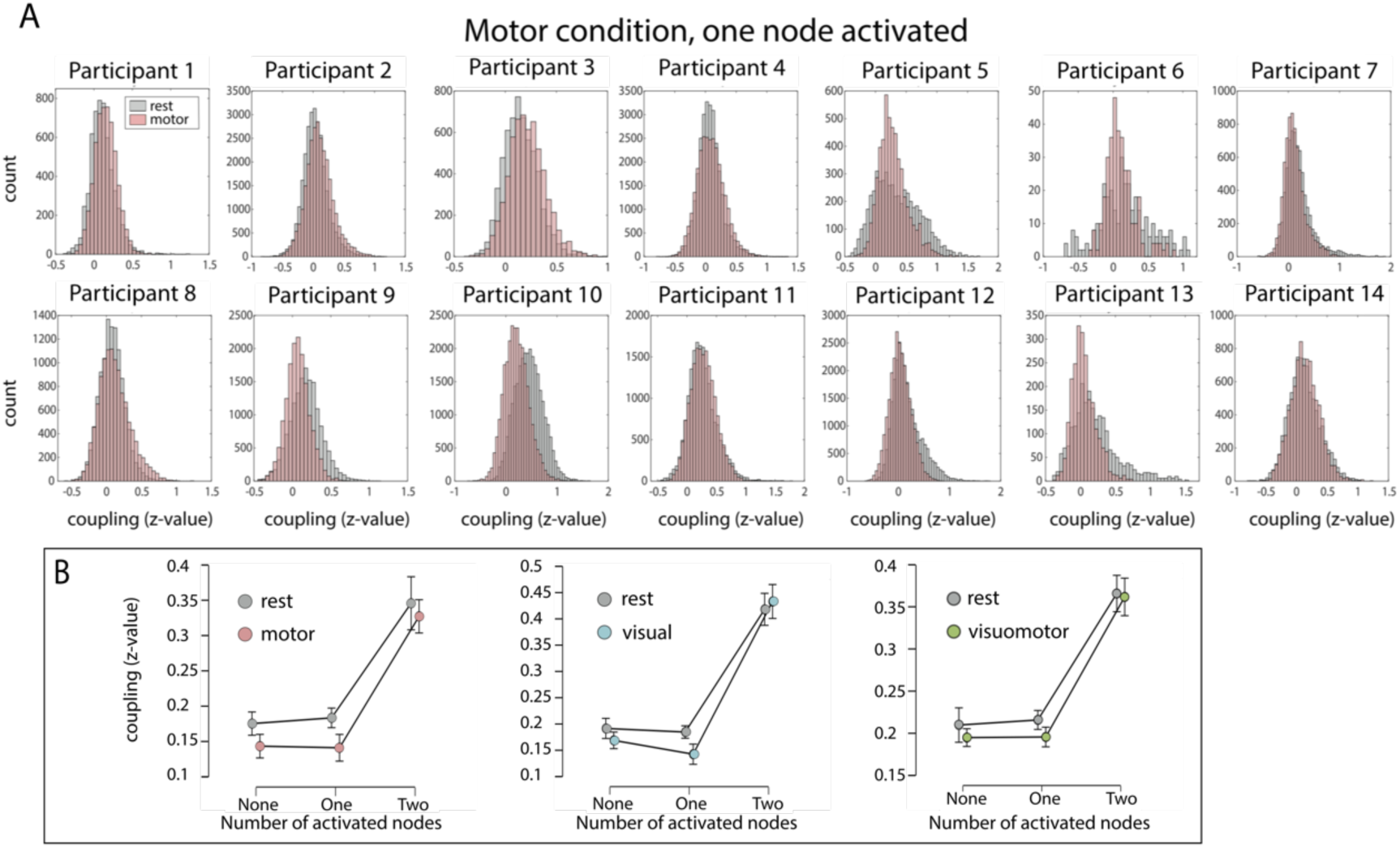
Relationship between task-evoked activation and differences between task-based and resting state FC. (A) Fisher z-transformed correlation coefficients between pairs of nodes, where only one of the nodes is activated during the motor steady-state condition, are displayed separately for each participant in the form of a histogram (yaxis depicting the number of node pairs in each bin). If intrinsic connectivity changes with task activation, then FC should consistently decrease in the task-state, when only one of the nodes is activated, as compared to the resting-state. Although this is true for some participants (e.g. Participants 9 and 10), others show an opposite trend (connectivity increased in the motor condition, see e.g. Participants 1-3). (B) Relationship between number of activated nodes and mean FC for the three task comparisons (motor – pink, visual – blue, visuomotor green). Note that while a significant main effect of the factor “number of activated nodes” (x-axis) can be observed, no significant interaction between activated nodes and task was found. Error bars indicate standard error of the mean.

### Correlations across intrinsic connectivity networks

Individual steady-state scans were temporally concatenated for each task condition (rest, motor, visual, visuomotor) to create task-specific 4D datasets. Each of the datasets was decomposed to 50 independent components using MELODIC (Multivariate Exploratory Linear Optimized Decomposition into Independent Components, part of FSL software). Group-level MELODIC output was matched, using spatial cross-correlations, to 10 canonical networks (Smith et al. 2009) obtained from the resting-state HCP data. The components with the highest correlation values from each task-specific MELODIC output were selected in a winner-takes all paradigm, to obtain 10 networks of interest independently for each of the 4 steady-state conditions (Fig. 3A, Supp. Fig. 9-12).

**Fig. 3.**
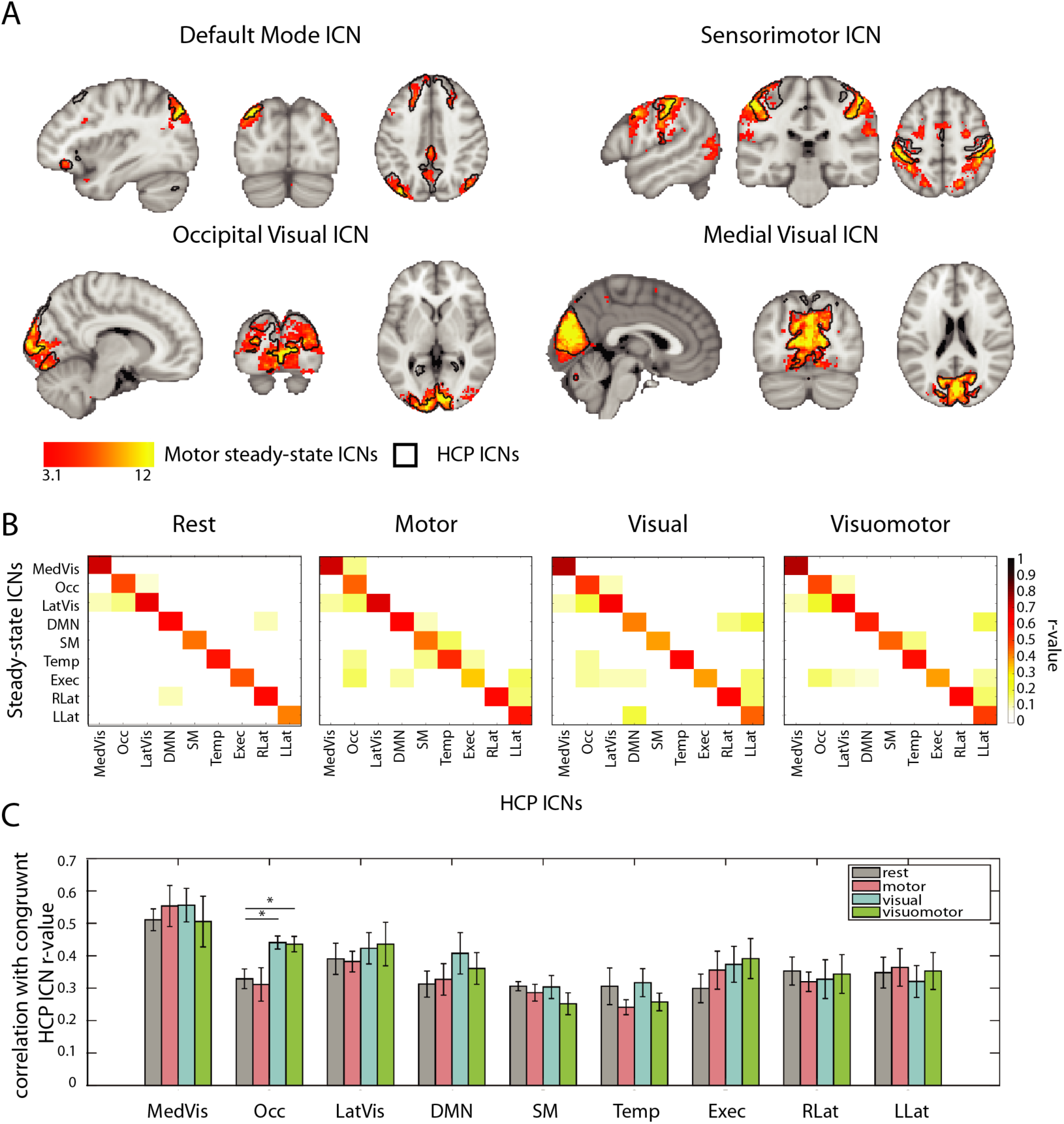
Spatial variability of ICNs between different steady-states. (A) Spatial maps of four task-related ICNs extracted from the motor condition (depicted in red-yellow scale) overlaid on the same ICNs extracted from the HCP data (depicted as black contours). (B) Whole brain correlation matrices of 9 major ICNs from the HCP data and their counterparts found in the steady-states data. Each ICN is correlated with all other ICNs. (C) Bar graph depicting mean spatial correlation coefficients (calculated from 100 bootstrapped ICA decomposition) of 9 major ICNs extracted from each of the steady-states conditions to their HCP counterparts. Note that only the Occipital Visual ICN (Occ) shows significant differences in its spatial correspondence to the HCP’s Occipital Network. Asterisks denote significance as determined using bootstrap percentile confidence intervals (see Methods). MedVis stands for Medial Visual ICN, Occ – Occipital Visual ICN, LatVis – Lateral Visual ICN, DMN – Default Mode Network, SM – Sensorimotor ICN, Temp – Temporal ICN, Exec – Executive ICN, RLat – Right Lateral ICN, LLat – Left Lateral ICN.

Upon inspection, the steady-state networks corresponding to the HCP cerebellum network were spatially diffuse in all four conditions, with its mean spatial correlation strength to the canonical cerebellum network being always under 0.2. The cerebellum network was therefore discarded from further analysis. Within each condition, the remaining 9 components were correlated against the HCP-derived networks across the entire brain (Fig. 3B).

Next, for each steady-states condition, we investigated the spatial consistency of the 9 identified networks with the HCP resting-state networks. To this extend, the individual participants’ datasets were combined into surrogate group fMRI datasets using a bootstrapping procedure (i.e. random sampling with replacement). For each steady-state condition 100 surrogate datasets per condition were created (Damoiseaux et al. 2006). Each surrogate dataset contained data from ten randomly selected participants, drawn from the given steady-state. For each surrogate dataset, 50 independent components were extracted using MELODIC and based on the spatial cross-correlations matched to the 10 canonical networks (Smith et al. 2009) obtained from the HCP’s data, resulting in 100 maps per condition for each of the 9 networks of interest. The obtained maps were spatially cross-correlated with the HCP-derived networks and for each network and each condition a bootstrapped sampling distribution of the r-values was built (Fig. 3C).

To quantify the differences in spatial correspondence of rest- and task-based ICNs to the resting-state HCP-derived networks, we built bootstrapped distributions of difference between r-values corresponding to the rest and motor; rest and visual; and rest and visuomotor steady-states. For each of those difference distributions, r-values were normalized using Fisher’s z-transform and bootstrap percentile confidence intervals were calculated. Confidence intervals not overlapping with zero indicated significant difference between networks derived from rest and a given task condition in their spatial correspondence to the canonical ICNs.

### Dual regression analysis

Finally, we quantified the differences in mean connectivity within the ICNs for each of the four steady-state conditions based on voxel-wise measures. We focused our analysis on four networks that overlapped with task-related activation changes during at least one of the employed steady-state conditions, as observed in the localizer: Default Mode Network ICN, Sensorimotor ICN, Occipital Visual ICN and Medial Visual ICN, as derived from the HCP dataset (Fig. 3A). To uncover network-specific functional connectivity differences we used the dual regression approach (Filippini et al. 2009). The main regressor of interest was the averaged time-course underlying the resting-state HCP component of interest, derived from each individual participant’s steady-state time-course, with individual voxels weighted based on their contribution to the group IC. The weighted ICA time-courses of the remaining HCP components of interest were also calculated within each individual participant, using the same procedure, and included as regressors of no interest, in a voxel-wise first-level GLM. The output values of this analysis are voxel-wise beta values, for each individual participant and in each steady-state, representing the strength of connectivity with the HCP components of interest, after accounting for partial contribution of all other ICs, for each individual participant and in each steady-state. For group statistical analysis, these beta maps were compared between the steady-state conditions using a 2 (motor task on/off) by 2 (visual task on/off) ANOVA design. This analysis was carried out using FSL randomize non-parametric permutation testing, with 5000 permutations, using a threshold-free cluster enhancement approach (Smith and Nichols 2009). This analysis was repeated independently for each of the four components of interest (Default Mode Network ICN, Sensorimotor ICN, Occipital Visual ICN and Medial Visual ICN), to identify the strength of connectivity between individual voxels and the given network. Since none of the interactions were significant, the analysis was restricted to two main effects per network of interest, resulting in 8 comparisons. To account for multiple comparisons, we adjusted the alpha value to 0.00625 (Bonferroni correction, see Fig. 4 for results).

**Fig. 4.**
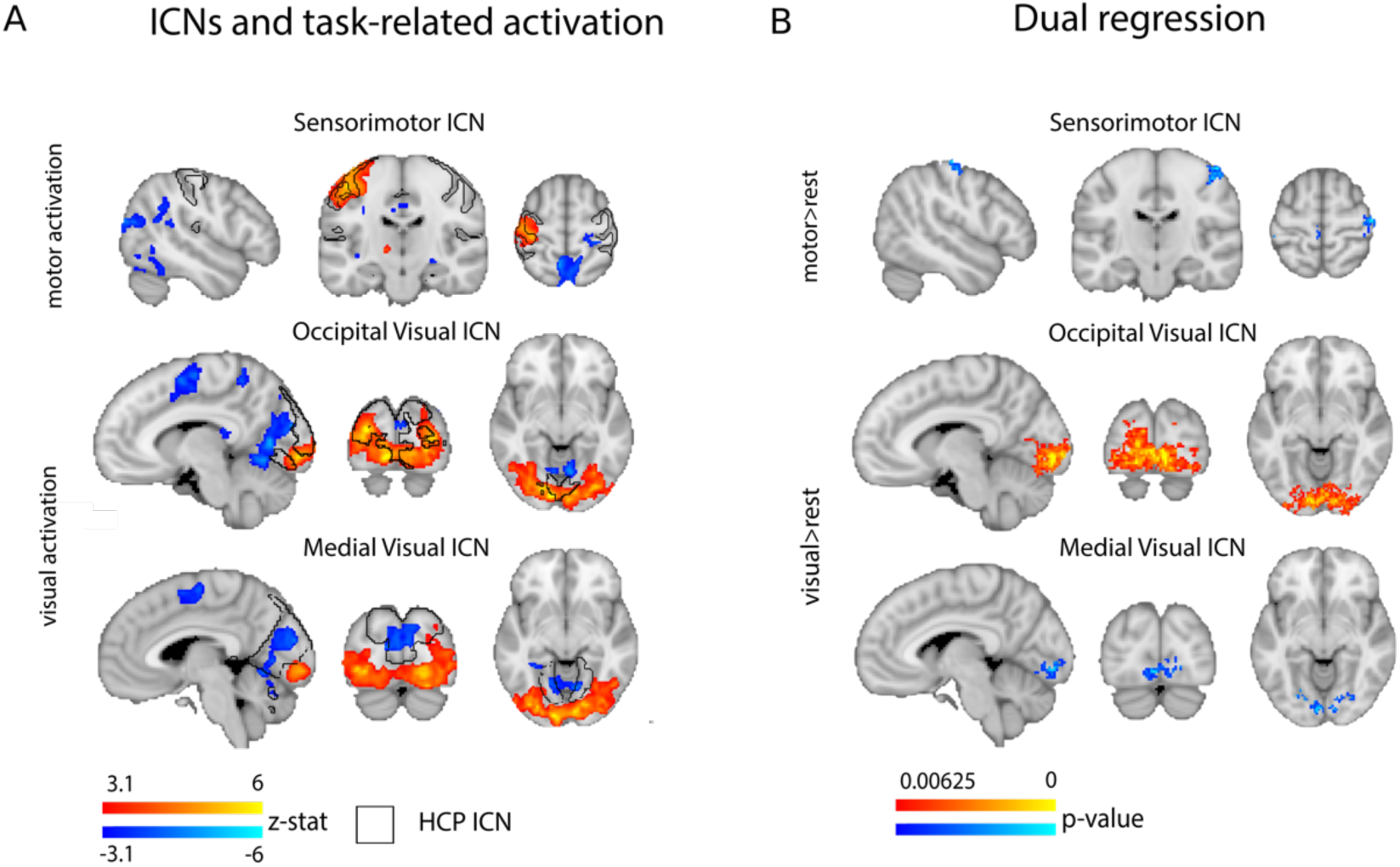
Intra-network FC differences between task and rest. (A) Brain regions activated by the motor condition overlap with the Sensorimotor ICN; brain regions activated during the visual condition overlap with the Occipital Visual ICN; and, brain regions deactivated during the visual condition overlap with the Medial Visual ICN. Brain (de)activation is shown in blue and red, the boundaries of the ICNs is illustrated by the black contour line. (B) Results of the dual regression analysis reveal: lower FC to the ipsilateral part of the Sensorimotor ICN during motor task; increased FC within the Occipital Visual ICN during visual task; and decreased connectivity between Medial Visual ICN and Occipital Visual ICN during the visual task.

## Results

### Task localizer

To evaluate changes in mean BOLD signal amplitude induced by each of the steady-state conditions (activation), we first examined group contrast maps for each of the task conditions versus rest, derived from the block-design localizer scan. As shown in Figure 1B, the visual and motor conditions resulted in the canonical visual/motor fMRI activation patterns (Allison et al. 2000). Since the finger-tapping motor task was performed using the right hand, during the motor condition positive activation could be observed in the left primary motor and pre-motor cortices (overlapping the Sensorimotor ICN), right cerebellum and the posterior part of the left putamen. As such, the activation profile only engaged a part of the canonical Sensorimotor ICN. In this condition, visual areas in the occipital cortex were associated with negative BOLD modulation (hereafter deactivation). The motor condition also resulted in deactivation, partially overlapping with areas formally known as comprising posterior regions of the Default Mode Network (e.g. posterior cingulate cortex, precuneus). The natural visual condition, involving abstract videos, activated low-level (foveal) and mid-level visual areas in the occipital and occipitotemporal cortex (overlapping with the Occipital Visual ICN) as well as higher-level visual areas in the occipitotemporal cortex (e.g. V4, hMT). This visual condition also resulted in deactivation in the low-level (peripheral) medial visual area (overlapping with the Medial Visual ICN) and in supplementary motor area. The combined visuomotor condition, involving both abstract videos and right-hand finger tapping, activated similar motor and visual areas as described for each condition separately, thereby providing an opportunity to study the combined activation impact across the individual visual and motor conditions. A comparison of the average time-course in the task-based and resting steady-state scans confirmed that the areas activated during the localizer were also activated during the steady state (motor vs. rest: t_(13)_=6.659; p<0.0001; visual vs. rest: t_(13)_=3.906; p=0.0018; visuomotor vs. rest: t_(13)_=2.159; p=0.05). The localizer data will be made available online at the Open Science Framework upon publication of the manuscript.

### Functional connectivity between nodes is not directly associated with changes in BOLD activation

This analysis aimed to determine whether changes in the intrinsic connectivity between task-related and task-unrelated nodes are associated with task-induced changes in mean fMRI activation across nodes. To examine the large-scale relationship between activation and connectivity, we parcellated the brain of each individual participant into functional nodes (see Methods). For each of the conditions (rest, motor, visual, visuomotor), we calculated a pairwise correlation between these nodes’ time-courses, which yielded a connectivity matrix. For each task-condition, pairs of nodes were then sorted based on whether both nodes, one node or neither of the nodes in each pair were positively activated during the visual, motor or visuomotor task localizer. The correlation values were Fisher Z-transformed and displayed in a histogram, separately for each of the participants (See Fig. 2A for motor condition, one node activated; see Supp. Fig. 1-8 for the other conditions).

If intrinsic FC is modulated by task demands (dependent on changes in node activation levels), then it should be increased/decreased in the task-state, when both/one of the nodes are activated by the task, respectively. This should result in a significant interaction between the number of nodes activated (zero, one, two) and the steady-state condition (task, rest). Conversely, we found that substantial proportion of FC changes is not significantly associated with activation. This was exemplified by a nonsignificant interaction between the number of activated nodes and the task (motor vs rest: F_(1.076,13.987)_=0.258, p=0.637; visual vs rest: F_(1.266,16.458)_=1.666, p=0.219; visuomotor vs rest: F_(1.083,14.074)_=0.357, p=0.576). The Bayes Factor (BF) for the interaction was below 0.33 for all three conditions (motor vs rest BF: 0.285, visual vs rest BF: 0.282, visuomotor vs rest BF: 0.182; all versus rest), providing positive evidence in favor of the null hypothesis (no interaction; Dienes 2014; Wetzels et al. 2011). We also found a significant main effect of the number of activated nodes in a pair (zero, one or two) on FC (Fig. 2B), indicating that nodes that usually activate together tend to show increased connectivity independent of the steady state (motor vs rest: F_(1.268,16.485)_=44.643, p<0.001; visual vs rest: F_(1.07,13.908)_=50.267, p<0.001; visuomotor vs rest: F_(1.131,14.703)_=46.374, p<0.001). Finally, we found no significant main effect of task, showing that connectivity was not significantly different between task- and rest-states (motor vs rest: F_(1,13)_=1.327, p=0.27; visual vs rest: F_(1,13)_=1.1688, p=0.299; visuomotor vs rest: F_(1,13)_=0.477, p=0.502). These null results were further examined using Bayesian statistics, where a threshold of Bayes factor (BF) < 1/3 was taken as positive (substantial) evidence in favor of the null hypothesis (no differences across networks; Kass & Raftery 1995; Wetzels *et al.* 2011). We found that the null hypothesis was supported for the visual (BF=0.272) and visuomotor conditions (BF=0.193), whereas the differences between the motor and rest conditions were ambiguous (BF=0.46).

### Spatial Consistency of ICNs across steady-state tasks

In the node analysis described above, we examined the local relationship between task-induced activation and FC. Although we found that the average FC between nodes was not reliably dependent on the activation changes between those nodes, it is possible that subtler changes in connectivity within each nodes category, impacting the overall spatial distribution of the intrinsic networks, were left undetected.

In our next analysis, we thus aimed to examine the stability of the ICNs across different steady-state conditions. In other words, we investigated whether the spatial patterns of the resting-state connectivity networks correspond with those found during task states. For this purpose, we decomposed each of the steady-state datasets to identify 9 canonical networks of interests, based on the resting-state HCP dataset (see Methods). All of the canonical ICNs were found in the ICA decomposition of each of the steady-state datasets. Moreover, spatial maps of both rest- and task-state networks showed high levels of consistency with the HCP-derived resting-state ICNs (see Fig. 3A for task-relevant networks resolved from motor steady-state; see Supp. Fig. 9-12 for all of the networks and conditions). Spatial correlations between congruent networks of the HCP and steady-state tasks were found to be considerably stronger than the correlations between incongruent networks (Fig. 3B) (average r-value for the intra-network correlations: 0.54-0.55, average r-value for the internetwork correlations: 0.01-0.02), demonstrating that the spatial distribution of the networks is largely preserved across task-states.

To quantify the spatial overlap of the data-driven ICNs across steady-states with their HCP-counterpart, we ran a bootstrapping analysis (see Methods, Fig. 3C), allowing us to quantify confidence intervals of spatial correlation values. In addition, this analysis provides an important opportunity to validate the quality of our data, considering our sample size was much smaller than that of the HCP. Despite the fact that HCP-derived ICNs were acquired during resting state while the task-based steady-states were acquired during activation of a range of brain areas (see Task localizer results), all of the networks showed similar levels of spatial overlap with the congruent HCP-networks. The exception was the Occipital Visual network which was more strongly correlated with its HCP-counterpart during visually related conditions (visual and visuomotor) than during rest (visual-rest difference score CI:-0.2173 to −0.0469, visuomotor-rest difference score CI: −0.2170 to −0.0464). Together these findings show that all ICNs, not only those activated by the experimental tasks (Smith et al. 2009), are remarkably preserved across steady-states, despite the changes in brain activation.

### Local differences in connectivity profiles

While our previous analysis showed that the ICNs are stable across different steady-state conditions, it is still possible that the task demands have a more localized impact on FC, which is insufficient to disrupt the ICNs global stability. It has previously been suggested that activation-driven changes in FC during a task-based block design are masked by the network’s intrinsic connectivity properties (Cole et al. 2014; Xie et al. 2017). To uncover potentially more subtle differences across the connectivity profiles of task-relevant networks, we employed a dual regression analysis. This analysis allowed us to look at differences in voxel-wise connectivity strengths within the networks of interest in each steady-state task condition. Importantly, unlike commonly used seed-based FC analyses, this approach allowed us to characterize the unique distribution of each of the networks while accounting for variability shared with other networks. Here we focused on four main networks, most relevant for the tasks we used (i.e. spatially overlapping with task-related activation changes, as found in the task-localizer; see Fig. 1B): The Medial and Occipital Visual networks, the Sensorimotor network and the Default Mode Network. We took advantage of our parametric design (visual activation on/off, motor activation on/off) to calculate one 2×2 ANOVA for each network. As no significant interactions were identified, we focused our analysis on the two main effects (see Methods). The Occipital Visual network overlapped with areas that were activated during the visual and visuomotor tasks. Accordingly, we found that the areas within the Occipital Visual network showed increase intra-network connectivity during those tasks, as compared to rest (Fig. 4). In other words, the Occipital Visual network becomes more strongly connected to itself in the visual conditions, which may potentially underlie its stronger correspondence to its HCP-derived counterpart during visual and visuomotor steady-states (Fig. 3C). The Medial Visual network overlapped with brain areas that were deactivated during the visual task. Accordingly, areas of this network showed reduced connectivity to the Occipital Visual ICN, which was activated during the visual conditions (Fig. 4). Finally, our motor tasks required the movement of only one hand, resulting in unilateral activation in the sensorimotor hand area contralateral to the task-related hand. Though the bilateral hand areas are typically coupled during resting state (Hahamy et al. 2015b) the inactive hand (ipsilateral) area showed decreased connectivity with the sensorimotor network during the motor conditions. No other significant results were found for other contrasts and networks under the adjusted threshold (alpha<0.00625). Note that similar results were also found under higher attentional load in the visuomotor condition (see Supporting Information), with additional FC modulations present in the Executive network. As such, it seems that areas that are activated/deactivated during the task may show increases/decreases in network coupling during task compared to rest, although these changes are contained within the relevant intrinsic connectivity networks and may thus be attributed to changes in the amplitude of variance of the driving signal (Duff et al. 2017; Nir et al. 2006).

## 4. Discussion

In the present study, we investigated the effects of regional steady-state brain activation on FC, by comparing resting-state FC measurements to steady-state task FC measurements. We used fMRI data from four steady-state tasks (Fig. 1A) to investigate modulations of FC due to changing task demands. In our first analysis, we looked at whole-brain changes in connectivity strength based on subject-specific activation profiles across nodes (Fig. 2). Here we benefitted from using naturalistic visual and motor tasks that result in robust and consistent activation in sensorimotor cortex. For example, a similar motor task has been recently shown to produce the largest effect sizes for changes in BOLD activation, as compared to cognitive tasks (Poldrack et al. 2017). Despite the fact that the activation was increased in a synchronized manner across task-specific nodes, we found no large-scale relation between FC and activation, suggesting that most FC changes may be better explained by network affiliation. To look at changes in the spatial attributes of FC we employed a data-driven approach based on ICA decomposition. We found that the resulting ICNs remain mostly undiminished during motor, visual, and visuomotor task conditions (Fig. 3). This was also the case when we examined an additional task, designed to increase attentional load and to better integrate across the visual and motor conditions (Supp. Fig. 14). This analysis demonstrates that the ICNs are not specific to resting state only but rather reflect the general state of functional brain organization. This observation is in accordance with previous studies postulating continuous intrinsic activity (Fox and Raichle 2007; Sadaghiani et al. 2010; Smith et al. 2009) and provides additional support for the idea that ICNs remain coupled within themselves even during task conditions (Calhoun et al. 2008; Greicius and Menon 2004; Hampson et al. 2006). Importantly, we show that even those networks that are partially (or not at all) activated by the task largely maintain their functional integrity during the steady-states. Together, our findings suggest that functional brain structure is defined by a set of stable intrinsic networks that are present across both low-level tasks and rest.

A number of previous studies have shown that FC of ICNs can still be measured during both sensorimotor and cognitive tasks in addition to rest (Calhoun et al. 2004; Calhoun et al. 2008; Moeller et al. 2009; Shirer et al. 2012; Smith et al. 2009; Sun et al. 2004). For instance, Sun et al. (2004) seeded certain areas of the motor cortex during a bimanual motor task and found that they could reproduce a FC pattern similar to the sensorimotor ICN seen during rest. Similarly, in macaques, Moeller et al. (2009) showed that independent component analyses of fMRI data acquired during movie-watching, rest and various visual tasks revealed FC networks that were highly similar across conditions. However, note that in these examples, the task-activation pattern corresponded with the spatial properties of the related ICNs. In our experimental design, we varied the extent of activation within and across ICNs: the visual task, comprised of colorful and slowly moving shapes, was designed to activate the entire Occipital Visual network; the motor task, comprised of unilateral hand movements, was designed to only activate parts of the bilateral sensorimotor ICN; the combined visual and motor condition was designed to evoke a summation of visual and motor activations (an effect previously observed by Calhoun et al. 2004), providing opportunity to observe inter-network interactions. Despite this diversified experimental design, we found that the ICNs were largely invariant to changed activation (see below for a discussion of induced task-changes, as identified in our final analysis). The exception to this rule was the Occipital Visual ICN, where spatial changes in the connectivity pattern were identified in the visual task conditions, compared to non-visual conditions (Fig. 3C). Our findings therefore demonstrate that the ICNs are robust to change, at least due to low task demands. This observation is consistent with recent evidence from cognitive tasks, which have been shown to introduce little variability to the gross structure of the FC networks (Gratton et al. 2018). Note however that other studies should determine whether these results can be replicated in a range of other steady-state tasks and paradigms, e.g. while activating only a proportion of the visual network and in tasks involving higher cognitive loads and/or fine motor precision.

One of the most prominent explanations of the immutability of the ICNs across task conditions is that the resting state fluctuations are stable and linearly superimposed on the task activation, as first postulated by e.g. Arieli (1996) and Fox et al. (2005). More recent studies (Cole et al. 2014; Gratton et al. 2016; Kimm et al. 2017; Xie et al. 2017) suggested that the functional brain architecture during both rest and task performance is dominated by the ICNs that are superimposed on any potential task-evoked FC changes. In other words, task-evoked FC changes occur in the presence of an intrinsic functional network architecture that extends across many or all brain states. Thus, the activation-driven changes can only be resolved after removing the intrinsic connectivity components from the data (Kim et al. 2017; Xie et al. 2017 though see also Dvir et al. 2017). Indeed, using the dual regression procedure, we found significant differences in connectivity profiles arising from different steady-state tasks. In summary, we found that some brain areas activated by the task (i.e. within the Occipital Visual ICN during visual task) tended to become more connected to the network. Areas that become deactivated by a task (Medial Visual ICN in visual, viusomotor and attention tasks; ipsilateral hand area in motor, visuomotor and attention tasks) become decoupled from the activated network. The observed decrease in FC within the motor system was previously reported by Shah et al. (2016) and Morgan and Price (2004). These authors hypothesized that this decrease in FC can be caused by the increased noise in the signal, induced by the finger tapping task. However, as those FC changes involve mainly the ipsilateral hemisphere (which does not activate during the task) rather than the entire sensorimotor network, we believe that the observed FC decrease originates from the lateralized activation characteristic of finger tapping (though we note that under higher attentional load the suppression was more extensive, see Supp. Fig. 14). All reported FC differences were, however, relatively localized to the activated/deactivated networks, suggesting that activation changes due to task demands only affect local connectivity within the network (as shown in Fig. 4). Exploring the linearity of the addition of the visual task to the motor task, we saw that there was relatively little interaction between the two conditions with respect to changes in connectivity, despite the fact that activation has been induced across both networks (see Supp. Fig. 15 for inter-network changes in the attention task). These findings resonate with our previous conclusion, that network affiliation may be the most important aspect of functional connectivity. Indeed, a likely framework for explaining the differences in the ICN’s task-specific connectivity profile may be the changes in the amplitude of the driving signal (Duff et al. 2017). Regardless, although relatively small, those connectivity changes are functionally meaningful and can potentially be used to distinguish between different cognitive tasks (Shirer et al. 2012) and participants (Tavor et al. 2016).

Despite widely established correspondence in rest-task network topography, recent studies have emphasized differences in FC patterns evoked by resting and task states (Betti et al. 2013; Buckner et al. 2013; Kim et al. 2017; Mennes et al. 2013). A common characteristic of most of the studies looking at activation-based changes in FC is that they use a block design and base their analyses on Psychophysiological Interactions (PPI). This procedure is based on the assumption that the global FC patterns can be initiated and stabilized within an order of seconds. However, it has been demonstrated that the characteristics of the spontaneous fluctuations change with time (Duff et al. 2008; Hutchison et al. 2013; see also Beckmann et al. 2005; Biswal et al. 1995; Cordes et al. 2000 for considerations of the FC frequency band and Hutchison et al. 2013 for dynamic functional connectivity). Here, we offer a paradigm that can help to ameliorate those confounds by using steady-state designs in which cognitive state is expected to be constant over time. Steady-state scans have been shown to be less susceptible to confounding factors than the block designs (Hampson et al. 2006) and to produce more consistent FC results (Fair et al. 2007). In the current study, we thus employed a set of simple yet extensively studied motor and visual tasks, allowing us to examine the interaction between BOLD responses to a particular stimulus and FC changes. By varying the visual stimulus and moving fingers across each scan, our steady-state tasks were specifically designed to minimize effects of fMRI adaptation (also known as repetition suppression) which regardless, usually contributes to only a very small proportion of the BOLD signal (Grill-Spector et al. 2006). Although performed continuously over a period of 5 minutes, we have found the steady-state tasks to be a good approximation of the task-localizer, in terms of reliably activating the same brain regions. Overall, we confirm previous observations that the intrinsic network architecture appears to be a canonical (default) state of the human brain’s functional network (Harmelech and Malach 2013), with task demands having a small effect on this state when considered in terms of overall brain organization.

Our results provide two opportunities for methodological impact. First, we show that node-to-node correlations are insensitive to localized task-based changes in FC. Those changes were only significantly observed when utilizing a dual regression approach that effectively regresses out any contributions to the FC time-course that are shared by the other networks and conditions. This suggests that dual regression can be effectively used for unmasking local FC changes, providing alternative means to previously used approaches (e.g. inter-subject functional correlations, Kimm et al. 2017; Ren et al. 2017). Second, our findings suggest that steady-state designs can be used to study ICNs. Many studies have shown that motion is a major source of variability in FC studies that can lead to erroneous results when comparing groups of participants (Power et al. 2012; Satterthwaite et al. 2012; Van Dijk et al. 2012). Moreover, it has been reported that task-based scans are associated with less head motion than classical resting-state scans (Vanderwal et al. 2015; Huijbers et al. 2017). Despite these methodological benefits, task-based scans have been avoided in FC studies due to the assumption that intrinsic connectivity cannot be robustly measured under task conditions. Our data challenge this assumption: we show that the intrinsic connectivity structure dominates over task-evoked FC and is thus reliably present across multiple types of task-based brain states. Our findings therefore demonstrate that the intrinsic functional network structure can be reliably assessed, and compared between different populations, during various steady-states; and potentially even when different participants are engaged in different minimally demanding steady-state tasks. As suggested by Vanderwal et al. (2015), this finding can facilitate data collection as it lowers the chances of participants falling asleep and significantly reduces their head movements. Furthermore, whereas resting state scans are largely uncontrolled (the final FC results can be altered by uncontrollable activations), steady-states paradigms offer a greater level of cognitive and experimental control, which may help to reduce variability in results or circumvent other confounds.

## Acknowledgements

TRM was funded by a Sir Henry Dale Fellowship jointly funded by the Wellcome Trust and the Royal Society (grant number 104128/Z/14/Z) and an ERC Starting Grant (grant number 0032-2-289-6121). EPD receives funding from Support for the Sick Newborn and their Parents (SSNAP) Oxford. NF was funded by the NIHR Oxford Health Biomedical Research Centre. The Wellcome Centre for Integrative Neuroimaging is supported by core funding from the Wellcome Trust (203139/Z/16/Z). Data were provided [in part] by the Human Connectome Project, WU-Minn Consortium (Principal Investigators: David Van Essen and Kamil Ugurbil; 1U54MH091657) funded by the 16 NIH Institutes and Centers that support the NIH Blueprint for Neuroscience Research; and by the McDonnell Center for Systems Neuroscience at Washington University.

## Conflicts of interest

All authors declare no financial or non-financial conflicts of interest.

## REFERENCES

Allison, J.D., Meador, K.J., Loring, D.W., Figueroa, R.E., Wright, J.C., 2000. Functional MRI cerebral activation and deactivation during finger movement. Neurology 54, 135–142.

Arieli, A., Sterkin, A., Grinvald, A., Aertsen, A., 1996. Dynamics of ongoing activity: explanation of the large variability in evoked cortical responses. Science 273, 1868–1871.

Beckmann, C.F., DeLuca, M., Devlin, J.T., Smith, S.M., 2005. Investigations into resting-state connectivity using independent component analysis. Philos Trans R Soc Lond B Biol Sci 360, 1001–1013.

Betti, V., Della Penna, S., de Pasquale, F., Mantini, D., Marzetti, L., Romani, G.L., Corbetta, M., 2013. Natural scenes viewing alters the dynamics of functional connectivity in the human brain. Neuron 79, 782–797.

Bianciardi, M., Fukunaga, M., van Gelderen, P., de Zwart, J.A., Duyn, J.H., 2011. Negative BOLD-fMRI signals in large cerebral veins. J Cerebral Blood Flow and Metabolism 31, 401–412.

Biswal, B.B., Mennes, M., Zuo, X.N., Gohel, S., Kelly, C., Smith, S.M., Beckmann, C.F., Adelstein, J.S., Buckner, R.L., Colcombe, S., Dogonowski, A.M., Ernst, M., Fair, D., Hampson, M., Hoptman, M.J., Hyde, J.S., Kiviniemi, V.J., Kotter, R., Li, S.J., Lin, C.P., Lowe, M.J., Mackay, C., Madden, D.J., Madsen, K.H., Margulies, D.S., Mayberg, H.S., McMahon, K., Monk, C.S., Mostofsky, S.H., Nagel, B.J., Pekar, J.J., Peltier, S.J., Petersen, S.E., Riedl, V., Rombouts, S.A., Rypma, B., Schlaggar, B.L., Schmidt, S., Seidler, R.D., Siegle, G.J., Sorg, C., Teng, G.J., Veijola, J., Villringer, A., Walter, M., Wang, L., Weng, X.C., Whitfield-Gabrieli, S., Williamson, P., Windischberger, C., Zang, Y.F., Zhang, H.Y., Castellanos, F.X., Milham, M.P., 2010. Toward discovery science of human brain function. Proc Natl Acad Sci U S A 107, 4734–4739.

Biswal, B.B., Yetkin, F.Z., Haughton, V.M., Hyde, J.S., 1995. Functional Conectivity in the Motor Cortex of resting Human Brain Using Echo-Planar MRI. Magnetic Resonance in Medicine 34, 537–541.

Buckner, R.L., Krienen, F.M., Yeo, B.T., 2013. Opportunities and limitations of intrinsic functional connectivity MRI. Nat Neurosci 16, 832–837.

Calhoun, V.D., Adali, T., Pekar, J.J., 2004. A method for comparing group fMRI data using independent component analysis: application to visual, motor and visuomotor tasks. Magn Reson Imaging 22, 1181–1191.

Calhoun, V.D., Kiehl, K.A., Pearlson, G.D., 2008. Modulation of temporally coherent brain networks estimated using ICA at rest and during cognitive tasks. Hum Brain Mapp 29, 828–838.

Cole, M.W., Bassett, D.S., Power, J.D., Braver, T.S., Petersen, S.E., 2014. Intrinsic and task-evoked network architectures of the human brain. Neuron 83, 238–251.

Cordes, D., Haughton, V.M., Arfanakis, K., Wendt, G.J., Turski, P.A., Moritz, C.H., Quigley, M.A., Meyerand, M.E., 2000. Mapping Functionally Related Regions of Brain with Functional Connectivity MR Imaging. AJNR Am J Neuroradiol 21, 1636–1644.

Costa, L., Smith, J., Nichols, T., Cussens, J., Duff, E.P., Makin, T.R., 2015. Searching Multiregression Dynamic Models of Resting-State fMRI Networks Using Integer Programming. Bayesian Analysis 10, 441–478.

Damoiseaux, J.S., Rombouts, S.A.R.B., Barkhof, F., Scheltens, P., Stam, C.J., Smith, S.M., Beckmann, C.F., 2006. Consistent resting-state networks across healthy subjects. PNAS 103.

Dienes, Z., 2014. Using Bayes to get the most out of non-significant results. Front Psychol 5, 781.

Duff, E., Johnston, L.A., Xiong, J., Fox, P.T., Mareels, I., Egan, G.F., 2008. The power of spectral density analysis for mapping endogenous BOLD signal fluctuations. Hum Brain Mapp 29, 778–790.

Duff, E., Makin, T.R., Madugula, S., Smith, S.M., Woolrich, M.W., 2013. Utility of Partial Correlation for Characterising Brain Dynamics: MVPA-based Assessment of Regularisation and Network Selection. Pattern Recognition in Neuroimaging. IEEE, Philadelphia, PA, USA.

Duff, E., Makin, T.R., Smith, S.M., Woolrich, M.W., 2017. Disambiguating brain functional connectivity. bioRxiv.

Dvir, R.B., Grossman, S., Furman-Haran, E., Malach, R., 2017. Quenching of spontaneous fluctuations by attention in human visual cortex. Neuroimage.

Eippert, F., Kong, Y., Winkler, A.M., Andersson, J.L., Finsterbusch, J., Buchel, C., Brooks, J.C., Tracey, I., 2017. Investigating resting-state functional connectivity in the cervical spinal cord at 3T. Neuroimage 147, 589–601.

Fair, D., Dosenbach, N.U.F., Church, J.A., Cohen, A.L., Brahmbhatt, S., Miezin, F.M., Barcg, D.M., Raichle, M.E., Petersen, S.E., Schlaggar, B.L., 2007. Development of distinct control networks through segregation and integration. PNAS 104, 13507–13512.

Feinberg, D.A., Moeller, S., Smith, S.M., Auerbach, E., Ramanna, S., Gunther, M., Glasser, M.F., Miller, K.L., Ugurbil, K., Yacoub, E., 2010. Multiplexed echo planar imaging for sub-second whole brain FMRI and fast diffusion imaging. PLoS One 5, e15710.

Filippini, N., MacIntosh, B.J., Hough, M.G., Goodwin, G.M., Frisoni, G.B., Smith, S.M., Matthews, P.M., Beckmann, C.F., Mackay, C.E., 2009. Distinct patterns of brain activity in young carriers of the APOE-E4 allele. PNAS 106, 7209–7214.

Fox, M.D., Greicius, M., 2010. Clinical applications of resting state functional connectivity. Front Syst Neurosci 4, 19.

Fox, M.D., Raichle, M.E., 2007. Spontaneous fluctuations in brain activity observed with functional magnetic resonance imaging. Nat Rev Neurosci 8, 700–711.

Fox, M.D., Snyder, A.Z., Zacks, J.M., Raichle, M.E., 2006. Coherent spontaneous activity accounts for trial-to-trial variability in human evoked brain responses. Nat Neurosci 9, 23–25.

Gilaie-Dotan, S., Hahamy-Dubossarsky, A., Nir, Y., Berkovich-Ohana, A., Bentin, S., Malach, R., 2013. Resting state functional connectivity reflects abnormal task-activated patterns in a developmental object agnosic. Neuroimage 70, 189–198.

Glasser, M.F., Coalson, T.S., Robinson, E.C., Hacker, C.D., Harwell, J., Yacoub, E., Ugurbil, K., Andersson, J., Beckmann, C.F., Jenkinson, M., Smith, S.M., Van Essen, D.C., 2016. A multi-modal parcellation of human cerebral cortex. Nature 536, 171–178.

Glasser, M.F., Sotiropoulos, S.N., Wilson, J.A., Coalson, T.S., Fischl, B., Andersson, J.L., Xu, J., Jbabdi, S., Webster, M., Polimeni, J.R., Van Essen, D.C., Jenkinson, M., Consortium, W.U.-M.H., 2013. The minimal preprocessing pipelines for the Human Connectome Project. Neuroimage 80, 105–124.

Gratton, C., Laumann, T.O., Nielsen, A.N., Greene, D.J., Gordon, E.M., Gilmore, A.W., Nelson, S.M., Coalson, R.S., Snyder, A.Z., Schlaggar, B.L., Dosenbach, N.U.F., Petersen, S.E., 2018. Functional brain networks are dominated by stable group and individual factors, not cognitive or daily variation. Neuron 98, 439–452.

Greicius, M.D., Menon, V., 2004. Default-Mode Activity during a Passive Sensory Task: Uncoupled from Deactivation but Impacting Activation. Journal of Cognitive Neuroscience 16, 1484–1492.

Greve, D.N., Fischl, B., 2009. Accurate and robust brain image alignment using boundary-based registration. Neuroimage 48, 63–72.

Griffanti, L., Salimi-Khorshidi, G., Beckmann, C.F., Auerbach, E.J., Douaud, G., Sexton, C.E., Zsoldos, E., Ebmeier, K.P., Filippini, N., Mackay, C.E., Moeller, S., Xu, J., Yacoub, E., Baselli, G., Ugurbil, K., Miller, K.L., Smith, S.M., 2014. ICA-based artefact removal and accelerated fMRI acquisition for improved resting state network imaging. Neuroimage 95, 232–247.

Grill-Spector, K., Henson, R., Martin, A., 2006. Repetition and the brain: neural models of stimulus-specific effects. Trends Cogn Sci 10, 14–23.

Guerra-Carrillo, B., Mackey, A.P., Bunge, S.A., 2014. Resting-state fMRI: a window into human brain plasticity. Neuroscientist 20, 522–533.

Hahamy, A., Behrmann, M., Malach, R., 2015a. The idiosyncratic brain: distortion of spontaneous connectivity patterns in autism spectrum disorder. Nat Neurosci 18, 302–309.

Hahamy, A., Macdonald, S.N., van den Heiligenberg, F., Kieliba, P., Emir, U., Malach, R., Johansen-Berg, H., Brugger, P., Culham, J.C., Makin, T.R., 2017. Representation of Multiple Body Parts in the Missing-Hand Territory of Congenital One-Handers. Curr Biol 27, 1350–1355.

Hahamy, A., Sotiropoulos, S.N., Henderson Slater, D., Malach, R., Johansen-Berg, H., Makin, T.R., 2015b. Normalisation of brain connectivity through compensatory behaviour, despite congenital hand absence. Elife 4.

Hampson, M., Driesen, N.R., Skudlarski, P., Gore, J.C., Constable, R.T., 2006. Brain connectivity related to working memory performance. J Neurosci 26, 13338–13343.

Harmelech, T., Malach, R., 2013. Neurocognitive biases and the patterns of spontaneous correlations in the human cortex. Trends Cogn Sci 17, 606–615.

Hermundstad, A.M., Bassett, D.S., Brown, K.S., Aminoff, E.M., Clewet, D., Freeman, S., Frithsen, A., Johnson, A., Tipper, C.M., Miller, M.B., Grafton, S.T., Carlson, J.M., 2013. Structural foundations of resting-state and task-based functional connectivity in the human brain. PNAS 110, 6169–6174.

Hu, D., Huang, L., 2015. Negative hemodynamic response in the cortex: Evidence opposing neuronal deactivation revealed via optical imaging and electrophysiological recording. J Neurophysiol 114, 2152–2161.

Hutchison, R.M., Womelsdorf, T., Allen, E.A., Bandettini, P.A., Calhoun, V.D., Corbetta, M., Della Penna, S., Duyn, J.H., Glover, G.H., Gonzalez-Castillo, J., Handwerker, D.A., Keilholz, S., Kiviniemi, V., Leopold, D.A., de Pasquale, F., Sporns, O., Walter, M., Chang, C., 2013. Dynamic functional connectivity: promise, issues, and interpretations. Neuroimage 80, 360–378.

Jenkinson, M., Bannister, P., Brady, M., Smith, S.M., 2002. Improved Optimization for the Robust and Accurate Linear Registration and Motion Correction of Brain Images. Neuroimage 17, 825–841.

Jenkinson, M., Beckmann, C.F., Behrens, T.E., Woolrich, M.W., Smith, S.M., 2012. Fsl. Neuroimage 62, 782–790.

Kelly, C., Castellanos, F.X., 2014. Strengthening connections: functional connectivity and brain plasticity. Neuropsychol Rev 24, 63–76.

Kim, D., Kay, K., Shulman, G.L., Corbetta, M., 2017. A New Modular Brain Organization of the BOLD Signal during Natural Vision. Cerebral Cortex, 1–17.

Mennes, M., Kelly, C., Colcombe, S., Castellanos, F.X., Milham, M.P., 2013. The extrinsic and intrinsic functional architectures of the human brain are not equivalent. Cereb Cortex 23, 223–229.

Moeller, S., Nallasamy, N., Tsao, D.Y., Freiwald, W.A., 2009. Functional connectivity of the macaque brain across stimulus and arousal states. J Neurosci 29, 5897–590

Moeller, S., Yacoub, E., Olman, C.A., Auerbach, E., Strupp, J., Harel, N., Ugurbil, K., 2010. Multiband multislice GE-EPI at 7 tesla, with 16-fold acceleration using partial parallel imaging with application to high spatial and temporal whole-brain fMRI. Magn Reson Med 63, 1144–1153.

Morgan, V.L., Price, R.R., 2004. The effect of sensorimotor activation on functional connectivity mapping with MRI. Magn Reson Imaging 22, 1069–1075.

Nir, Y., Hasson, U., Levy, I., Yeshurun, Y., Malach, R., 2006. Widespread functional connectivity and fMRI fluctuations in human visual cortex in the absence of visual stimulation. Neuroimage 30, 1313–1324.

O’Reilly, J.X., Woolrich, M.W., Behrens, T.E., Smith, S.M., Johansen-Berg, H., 2012. Tools of the trade: psychophysiological interactions and functional connectivity. Soc Cogn Affect Neurosci 7, 604–609.

Poldrak, R.A., Baker, C.I., Durnez, J., Gorgolewski, K.J., Matthews, P.M., Munafo, M.R., Nochols, T.E., Poline, J.B., Vul, E., Yarkoni, T., 2017. Scanning the horizon: towards transparent and reproducible neuroimaging research. Nat Rev Neurosci 18, 115–126.

Power, J.D., Barnes, K.A., Snyder, A.Z., Schlaggar, B.L., Petersen, S.E., 2012. Spurious but systematic correlations in functional connectivity MRI networks arise from subject motion. Neuroimage 59, 2142–2154.

Ren, Y., Nguyen, V.T., Guo, L., Guo, C.C., 2017. Inter-subject Functional Correlation Reveal a Hierarchical Organization of Extrinsic and Intrinsic Systems in the Brain. Sci Rep 7, 10876.

Sadaghiani, S., Hesselmann, G., Friston, K.J., Kleinschmidt, A., 2010. The relation of ongoing brain activity, evoked neural responses, and cognition. Front Syst Neurosci 4, 20.

Sadaghiani, S., Kleinschmidt, A., 2013. Functional interactions between intrinsic brain activity and behavior. Neuroimage 80, 379–386.

Satterthwaite, T.D., Wolf, D.H., Loughead, J., Ruparel, K., Elliott, M.A., Hakonarson, H., Gur, R.C., Gur, R.E., 2012. Impact of in-scanner head motion on multiple measures of functional connectivity: relevance for studies of neurodevelopment in youth. Neuroimage 60, 623–632.

Shah, L.M., Cramer, J.A., Ferguson, M.A., Birn, R.M., Anderson, J.S., 2016. Reliability and reproducibility of individual differences in functional connectivity acquired during task and resting state. Brain Behav 6, e00456.

Shih, Y.Y., Chen, C.C., Shyu, B.C., Lin, Z.J., Chiang, Y.C., Jaw, F.S., Chen, Y.Y., Chang, C., 2009. A new scenario for negative functional magnetic resonance imaging signals: Endogenous neurotransmission. J Neurosci 29, 3036–3044.

Shirer, W.R., Ryali, S., Rykhlevskaia, E., Menon, V., Greicius, M.D., 2012.Decoding subject-driven cognitive states with whole-brain connectivity patterns. Cereb Cortex 22, 158–165.

Smith, S.M., 2002. Fast robust automated brain extraction. Hum Brain Mapp 17, 143–155.

Smith, S.M., 2004. Overview of fMRI analysis. Br J Radiol 77 Spec No 2, S167–175.

Smith, S.M., Nichols, T.E., 2009. Threshold-free cluster enhancement: addressing problems of smoothing, threshold dependence and localisation in cluster inference. Neuroimage 44, 83–98.

Smith, S.M., Fox, P.T., Miller, K.L., Glahn, D.C., Fox, P.M., Mackay, C.E., Filippini, N., Watkins, K.E., Toro, R., Laird, A.R., Beckmann, C.F., 2009. Correspondence of the brain’s functional architecture during activation and rest. PNAS 106.

Smith, S.M., Jenkinson, M., Woolrich, M.W., Beckmann, C.F., Behrens, T.E., Johansen-Berg, H., Bannister, P.R., De Luca, M., Drobnjak, I., Flitney, D.E., Niazy, R.K., Saunders, J., Vickers, J., Zhang, Y., De Stefano, N., Brady, J.M., Matthews, P.M., 2004. Advances in functional and structural MR image analysis and implementation as FSL. Neuroimage 23 Suppl 1, S208–219.

Smith, S.M., Andersson, J., Auerbach, E.J., Beckmann, C.F., Bijsterboch, J., Douaud, G., Duff, E., Feinberg, D.A., Griffanti, L., Harms, M.P., Kelly, M., Laumann, T., Miller, K.L., Moeller, S., Peterson, S., Power, J., Salimi-Khorshidi, G., Snyder, A.Z., Vu, A., Woolrich, M.W., Xu, J., Yacoub, E., Ugurbil, K., Van Essen, D., Glasser, M.F., 2013. Resting-state fMRI in Human Connectome Project. Neuroimage 80, 144–168.

Spadone, S., Della Penna, S., Sestieri, C., Betti, V., Tosoni, A., Perrucci, M.G., Romani, G.L., Corbetta, M., 2015. Dynamic reorganization of human resting-state networks during visuospatial attention. Proc Natl Acad Sci U S A 112, 8112–8117.

Sun, F.T., Miller, L.M., D’Esposito, M., 2004. Measuring interregional functional connectivity using coherence and partial coherence analyses of fMRI data. Neuroimage 21, 647–658.

Tavor, I., Parker Jones, O., Mars, R.B., Smith, S.M., Behrens, T.E., Jbabdi, S., 2016. Task-free MRI predicts individual differences in brain activity during task performance. Science 352, 216–2

The JASP Team, 2017. JASP Version 0.8.2.

Van Dijk, K.R., Hedden, T., Venkataraman, A., Evans, K.C., Lazar, S.W., Buckner, R.L., 2010. Intrinsic functional connectivity as a tool for human connectomics: theory, properties, and optimization. J Neurophysiol 103, 297–321.

Van Dijk, K.R., Sabuncu, M.R., Buckner, R.L., 2012. The influence of head motion on intrinsic functional connectivity MRI. Neuroimage 59, 431–438.

Wetzels, R., Matzke, D., Lee, M.D., Rouder, J.N., Iverson, G.J., Wagenmakers, E.J., 2011. Statistical Evidence in Experimental Psychology: An Empirical Comparison Using 855 t Tests. Perspect Psychol Sci 6, 291–298.

Wilf, M., Strappini, F., Golan, T., Hahamy, A., Harel, M., Malach, R., 2017. Spontaneously Emerging Patterns in Human Visual Cortex Reflect Responses to Naturalistic Sensory Stimuli. Cereb Cortex 27, 750–763.

Xie, H., Calhoun, V.D., Gonzalez-Castillo, J., Damaraju, E., Miller, R., Bandettini, P.A., Mitra, S., 2017. Whole-brain connectivity dynamics reflect both task-specific and individual-specific modulation: A multitask study. http://dx.doi.org/10.1016/j.neuroimage.2017.05.050

Zuo, N., Yang, Z., Liu, Y., Li, J., Jiang, T., 2-1 Both activated and less-activated regions identified by functional MRI reconfigure to support task executions. Brain Behav 8(1), e00893.

